# Quantum dot probes for the quantitative study of drug transport by the MacAB TolC efflux pump in lipid scaffolds

**DOI:** 10.1101/2020.06.16.154831

**Authors:** Hager Souabni, William Batista dos Santos, Quentin Cece, Dhenesh Puvanendran, Martin Picard

**Affiliations:** Université de Paris, Laboratoire de Biologie Physico-Chimique des Protéines Membranaires, CNRS UMR 7099, F-75005 Paris, France; Institut de Biologie Physico-Chimique, Fondation Edmond de Rothschild pour le développement de la recherche Scientifique, F-75005 Paris, France

## Abstract

ABC tripartite efflux pumps are macromolecular membrane protein machineries that expel a large variety of drugs and export virulence factors from Gram negative bacteria. Using a lipid scaffold mimicking the two-membrane environment of the transporter and designing spectroscopic conditions allowing the monitoring of both ATP hydrolysis and substrate transport in real time, we show that MacAB-TolC accommodates transport and energy consumption with high coupling efficiency.

## Introduction

The rapid emergence of bacteria capable to resist multiple antibiotics has led to the development of multidrug resistance (MDR), that frequently puts in danger the lives of patients across the world. In extreme cases, bacterial strains have been shown to develop resistance against almost all known chemotherapeutic agents and antibiotics. In Gram negative bacteria, resistance is mostly due to the combination of protein assemblies, called efflux pumps, that actively export noxious compounds, and of an impermeable double membrane barrier composed of lipids, sugars and peptidoglycans that oppose to the passive diffusion of molecules inside the cell^1^. Efflux pumps are composed of an inner membrane transporter, a periplasmic protein attached to the inner membrane, and an extrusion channel inserted in the outer membrane. They assemble to allow for an efficient transport of drugs, dyes or detergents outside the cell, bypassing the periplasm. The first line of defense is composed of efflux pumps that belong to the Resistance-Nodulation-Cell-Division (RND) superfamily, with, as most studied members AcrAB-TolC tripartite system from *E. coli* or MexAB OprM from *Pseudomonas aeruginosa*. Recently, the MacAB TolC tripartite efflux pump from *Escherichia coli* raised additional and increasing interest^2^. By contrast to the RND efflux pumps that perform transport by consuming protons from the periplasm, MacB is a membrane protein from the ABC superfamily that performs ATP-driven active translocation of substrates across the membrane. Substrate translocation occurs through ATP hydrolysis cycles concerted with conformational changes in the transmembrane domains where the substrate binds on one side of the membrane and is released to the other. A dimer of MacB assembles with an hexamer of MacA (a membrane fusion protein, MFP), and a trimer of TolC (multifunctional outer membrane channel protein). MacB was the first antibiotic-specific ABC drug exporter experimentally identified in a Gram negative bacterium^3^, initially shown to export macrolide compounds containing 14- and 15-membered lactones. Recently several structures described this complex with atomic detail in resting and drug transport states, suggesting that assembly and function of the pump are allosterically-coupled by a structural switch that synchronizes initial ligand binding and channel opening^4–8^. Biochemical studies showed that the ATPase activity of MacB is influenced by the presence of MacA and this was ascribed to its putative role in communicating the presence or absence of substrate in the periplasm to the nucleotide-binding and hydrolyzing domains (NBDs), located 100 Å away from the periplasm. Overall, questions regarding the modes of assembly and requisites for optimal function have been well-characterized^9–12^ but the puzzle of how exactly transport is coupled (*i.e* how much ATP it takes to complete a full transport cycle), is still a matter of debate and controversies. This question is central to the mechanic description of ABC transporters but is subjected to pitfalls because of the hydrophobic nature of most ABC substrates that may spontaneously partition into membranes in a non-protein fashion. As a consequence, kinetic measurements of both energy consumption and substrate transport are scarce in the literature^13–19^ and obviously such a study has never been achieved so far for an ABC tripartite transporter. We have decided to tackle the question of ATP:substrate coupling efficiency by the MacAB TolC efflux pump from *Escherichia coli* upon reconstitution of the proteins into lipid membranes and correlate ATP hydrolysis with transmembrane transport.

## Results

### Overall principle of the assay

Monitoring the activity of tripartite efflux pump from Gram negative bacteria is very challenging because they span the double membrane, and transport occurs from one environment to an other (see schematic representation Figure 1a). In the past, we have designed procedures to mimic efflux by reconstituting the respective protein partners in lipid vesicles and monitor *in vitro* transport through a reconstituted pump^20–23^. Following similar lines, we decided to set up a membrane-based system capable of mimicking the MacAB TolC tripartite complex. To that end, MacA and MacB were reconstituted into nanodiscs composed of POPC (1-palmitoyl-2-oleoyl-sn-glycero-3-phosphocholine) stabilized by a membrane scaffold protein (MSP, MSP1D1) that wraps around the hydrophobic core of the lipid discs (see supplementary section for details regarding materials and methods). Similarly, TolC was reconstituted into proteoliposomes composed of DOPC (1,2-dioleoyl-sn-glycero-3-phosphocholine) at a protein:lipid ratio such that, in average, one trimer of TolC is present per liposome. Tripartite complex formation was achieved by mixing MacAB nanodiscs and TolC-proteoliposomes in a 1:1 molar ratio (Figure 1b). Upon assembly, the MacAB nanodisc docks onto TolC, forming a complex capable to transport substrates from the buffering solution (representing the periplasmic space) to the interior of the proteoliposome (representing the bacterial outer medium), at the expense of ATP hydrolysis. As mandatory steps towards our goal to evaluate energy coupling, we have set up two independent procedures, described below, to monitor substrate transport (Figure 1c and Figure 2a, b) and ATP hydrolysis (see Figure 1d and Figure 2c) in real time.

**Figure 1.**
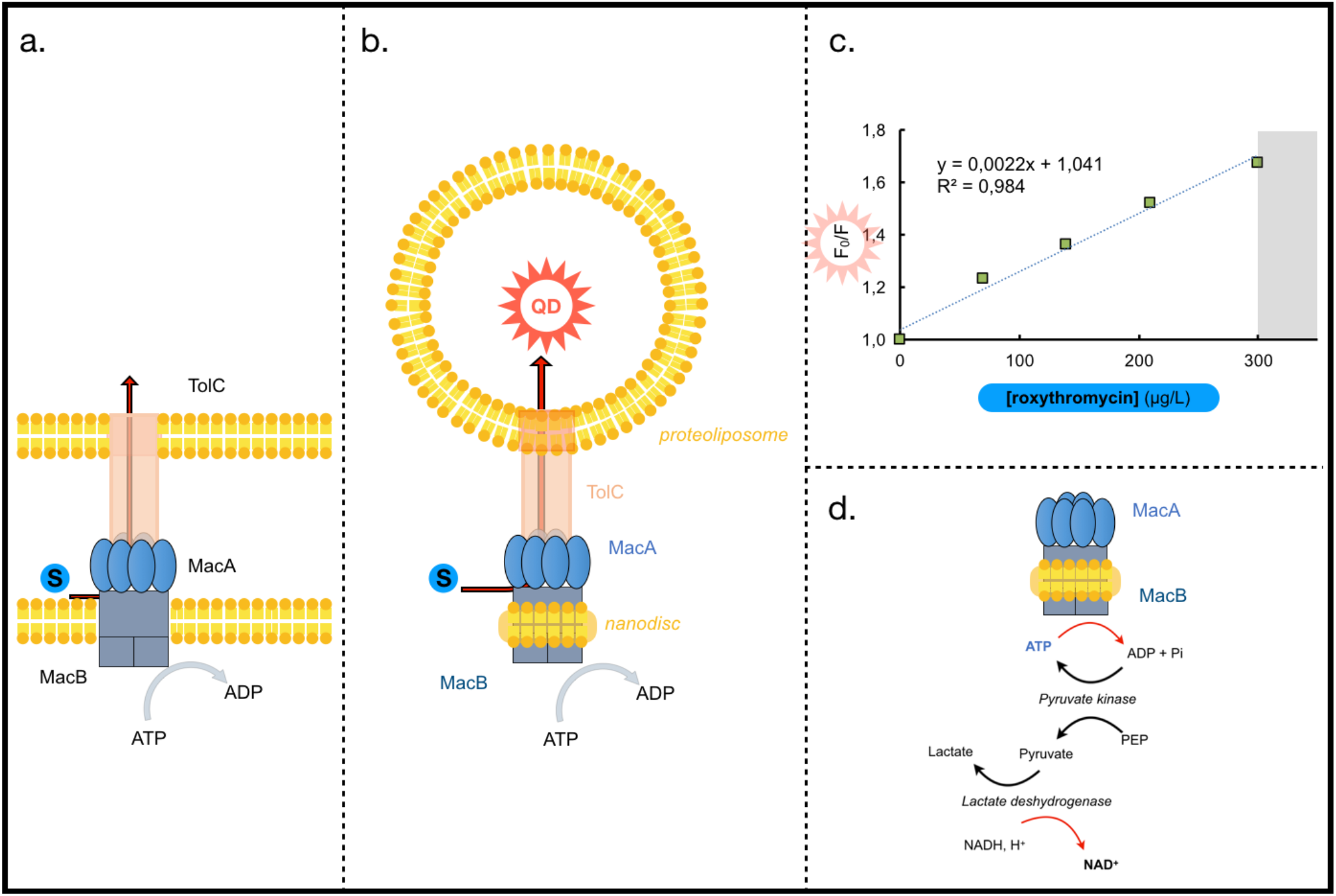
Rationale of the assay. *a. Schematic representation of the efflux pump assembly in E. coli membranes*. The MacAB-TolC pump encompasses the two-membrane barrier the Gram negative bacterium. A trimer of TolC, embedded in the outer membrane, protrudes into the periplasm and docks onto a hexamer of MacA that mediates the interaction with the periplasmic domain of MacB, which is a dimer. *b. Schematic representation of the in vitro assay*. MacA and MacB, in lipid nanodiscs and OprM in proteoliposomes form the tripartite complex upon mixing and incubation. Transport through the whole efflux pump is studied by monitoring ATP hydrolysis and antibiotic transport from the solution, representing the periplasmic space to the interior of the liposome representing the exterior of the bacteria. *c. Stern Volmer plot*. Quenching of the quantum dots fluorescence in the presence of increasing concentrations of roxithromycin. Results are best described by the Stern–Volmer type equation: F0/F = 1+ Ksv[S], where F and F0 are the fluorescent intensities of the QDs at a given roxithromycin concentration and in a roxithromycin-free solution, respectively. *d. NADH-coupled ATPase assay*. Hydrolysis of ATP is coupled to the oxidation of NADH as a consequence of two consecutive enzymatic reactions. First, ADP generated upon ATP hydrolysis is regenerated to ATP by pyruvate kinase (PK) that catalyzes the transition of phosphoenolpyruvate (PEP) to pyruvate. Pyruvate is then reduced to lactate by lactate dehydrogenase (LDH), which concomitantly catalyzes the oxidation of NADH. Hence, decrease in ATP concentration is directly and stoichiometrically correlated to the decrease in NADH concentration.

**Figure 2.**
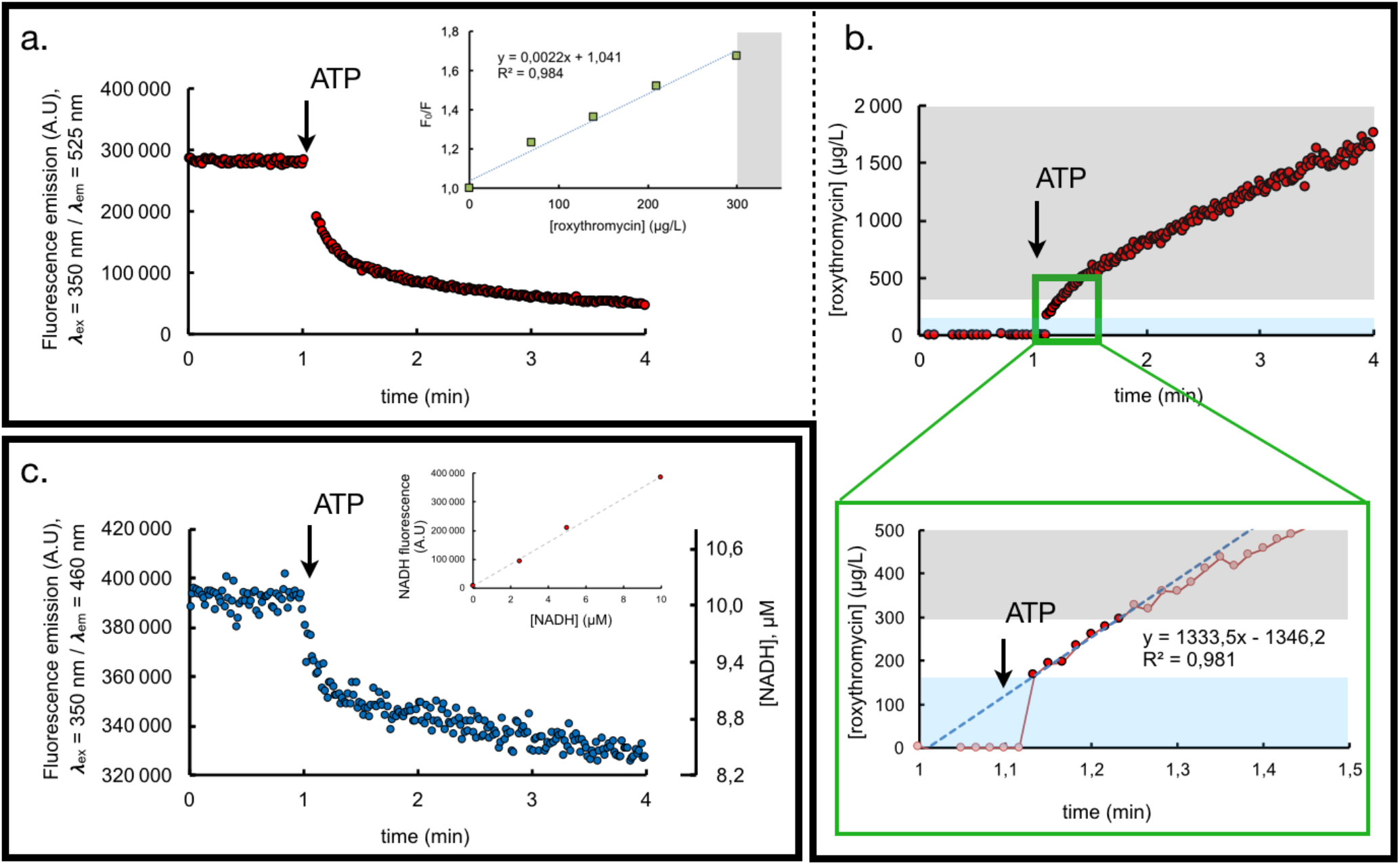
Real-time monitoring of roxithromycin transport (2a, b) and ATP hydrolysis (2c) within a reconstituted MacAB-TolC pump. a) CdTe quantum dot fluorescence is measured as a function of time after 10-fold dilution of the MacAB / TolC complex into 20 mM Tris pH 8, 50 mM NaCl, 2mM MgCl_2_, addition of 7μM roxithromycin (not shown here for the sake of clarity) and of 1 mM ATP (represented by the arrow). The MacAB nanodiscs (25μg MacB, as standardized by SDS PAGE densitometry, see supplemental Figure S2) and TolC proteoliposomes were preincubated for at least 1h at room temperature prior to their addition in the fluorescence cuvette, roxithromycin was added in the cuvette at least 10 minutes before addition of ATP. Changes due to dilution have been corrected for. *inset*: Quenching of the quantum dots fluorescence in the presence of increasing concentrations of roxithromycin shows that the fluorescence variation is proportional to the quantity of quencher up to a roxithromycin concentration of 300μg/L. b) Thanks to the above-described proportionality relationship, CdTe fluorescence quenching is converted into a quantity of roxithromycin transported as a function of time into the lumen of the liposome. The grey area represents the substrate concentration range for which proportionality does not hold. Values therein must not be taken into account. The green panel is a zoom of the region of interest. The blue area highlights the first fast phase of transport. c) NADH fluorescence is measured as a function of time after 10-fold dilution of the MacAB nanodiscs (*c.a* 10 μg MacB, as standardized by SDS PAGE densitometry, see supplemental Figure S1) into 20 mM Tris pH 8, 50 mM NaCl, 2mM MgCl_2_ containing the coupled-enzyme assay (see Methods for details) and addition of 1 mM ATP (represented by the arrow). Changes due to dilution have been corrected for. *inset*: linearity of NADH fluorescence over the range of concentration used in the assay. NADH fluorescence was measured with an excitation wavelength set at 350 nm and emission at 460 nm. Measurements were performed at room temperature (20°C) in 20 mM Tris pH 8, 50 mM NaCl, 2mM MgCl_2_. The proportionality coefficient allowed the conversion of the fluorescence changes (primary *y* axis, left) into [NADH] variations (secondary *y* axis, right).

### QD-based monitoring of roxithromycin

The main challenge of our study was to design a spectroscopic assay accounting for the realtime, quantitative, transport activity of the pump. To that end, we used quantum dots as fluorescent sensors for the presence of antibiotics. Quantum dots (QDs) are nanometer-sized semiconductor crystals that exhibit excellent brightness and superior photostability compared to conventional fluorophores^24^. They are ideal for applications that require high sensitivity, minimal interference with concurrent probes and long-term photostability. As additional asset, the fluorescence of CdTe quantum dots have been shown to be quenched in a dose-dependent manner in the presence of roxithromycin^25^, a semi-synthetic macrolide derived from erythromycin and known to be a very good substrate of the pump^5^. As a first control, we made sure that the range over which the concentration of analyte can be accurately monitored is compatible with our measurements. To that purpose, we measured QD fluorescence in the presence of increasing concentrations of roxithromycin. The ratio F0/F is plotted as a function of the quencher concentration (Stern–Volmer plot) to ascertain that we are dealing with a dynamic quenching (Figure 2a, inset) and the variation is found to be linear, with a slope corresponding to a Stern–Volmer constant in perfect agreement with that measured by Peng and collaborators^25^. From all the above, it seemed appealing to take advantage of the specific fluorescence quenching of roxithromycin on QD probes to detect and quantify the passage of the antibiotic into the TolC acceptor proteoliposomes upon transport through the tripartite reconstituted pump (Figure 1b). Indeed, when MacAB nanodiscs are preincubated with roxithromycin and TolC QD-entrapped proteoliposomes, we measure a steep and biphasic decrease of QD fluorescence as soon as ATP is added (Figure 2a). At steady state, we measure a rate of 1333 μg roxithromycin transported per mL and per minute (see Figure 2b), *i.e* 1.6 mM^-1^/min^-1^. Deconvoluted to the quantity of protein present in the tube and to the estimated average internal volume of a liposomes of 100 nm diameter size, we estimate the rate of transport to about 65 nmol.mg^-1^.min^-1^ (measurement performed in triplicate, see Supplementary Figure 3 for detail of the calculation). As mandatory controls, we also checked that no such quenching is observed when the experiment is performed in the presence of orthovanadate or in the absence of QD inside the TolC proteoliposome (see supplementary Figure 4b).

### ATP hydrolysis

Kinetic studies of ATPase activity can be spectroscopically monitored using a Nicotinamide adenine dinucleotide (NADH)-coupled ATPase assay (see Figure 1d) where decrease in ATP concentration is directly and stoichiometrically correlated to the decrease in NADH concentration^26^. Classically, the latter is deduced from variations in NADH specific absorbance at 340nm. However in our case, because MacB turnover number is known to be low (activities in the range of the tens of nmol.mg^-1^.min^-1^) this approach would require very high amounts of protein in order to reach ATP consumption levels sufficient for an observable decreasing of the absorbance signal. Attempting to measure ATP hydrolysis under these conditions gave rise to optical artifacts (because of light scattering artifacts) and rendered interpretation impossible. Considering the fact that in diluted systems, fluorescence emission is proportional to the concentration of the fluorophore, we made the assay more sensitive by monitoring ATP hydrolysis based on the variations of NADH intrinsic fluorescence. As a preliminary step, we have verified that indeed fluorescence is proportional to NADH over the concentration range used in our study (Figure 2c, inset). Measuring NADH fluorescence as a function of time in the coupled-enzyme assay for MacAB nanodiscs (Figure 2c) shows that ATP hydrolysis is biphasic, as previously shown by Lin *et al*.^12^. The specificity of the ATPase stimulation was further confirmed by using a MacB D169N mutant containing substitution in a conserved catalytic residue, involved in Mg^2+^ binding, proposed to play a central role in the ATP hydrolysis reaction by acting as a general base (Supplementary Material Figure 4a). When the slope of the fluorescence signals is now converted into ATPase activities, we find that, subsequent to the ATP hydrolysis “burst”, the steady state specific activity of MacAB (WT) is equal to 21,9 nmol.mg^-1^.min^-1^ (average of 9 independent measurements, in the presence or in the absence of substrate). This value is on the same order of magnitude as that previously shown for MacB pump^9,12^, and, similarly to the latter studies, we report that the ATPase activity is marginally affected by the presence of substrate.

## Discussion

Measurement of energy consumption is a rather straightforward and routine assay to determine the activity of ABC transporters but it is not always a perfectly relevant index because futile hydrolysis cycles can take place, leading to ATP hydrolysis rates that are not always stoichiometrically coupled to substrate transfer. For example, eukaryotic homodimeric TAPL complex undergoes ATP hydrolysis that are highly uncoupled from substrate transport (20-100 ATP hydrolyzed per translocated peptide) consistent with a high degree of futile hydrolysis cycles^14^ while the heterodimeric TAP complex shows a strict coupling between peptide transport and ATP hydrolysis^13^. Whether futile, wasteful, expenditure of ATP is an intrinsic feature of ABC transporters or an artefactual consequence of the manipulation of detergent-solubilized proteins is a matter of debate^27^. We show here that at steady state, MacAB-TolC transports 65 nmol roxithromycin per mg MacB and per minute at the expense of 20 nmol ATP hydrolyzed, meaning that, in our hands, MacAB is a very efficient transporter. Our results are in accordance with the general idea behind the “fireplace bellow” model proposed by Vassilis Koronakis for the same pump^4^. In the latter paper, it is postulated that ATP hydrolysis is not used to specifically transport the substrate across the membrane but instead to mechanically transmit conformational changes from the NBDs, in the cytoplasm where ATP hydrolysis takes place, to the large periplasmic domain, where transport occurs upon sequential opening and closing of a promiscuous cavity (see schematic representation Figure 3a). In other words, ATP is consumed at a constant pace, leading to the shrinkage of the periplasmic cavity and to the efflux of whatever molecule(s) present in the periplasmic pocket at that point of the catalytic cycle. This accounts for both the lack of substrate-induced activity (ATPase activity is of the same order of magnitude irrespective of the presence of substrate) and to the surprisingly large number of molecules transported per turnover.

**Figure 3.**
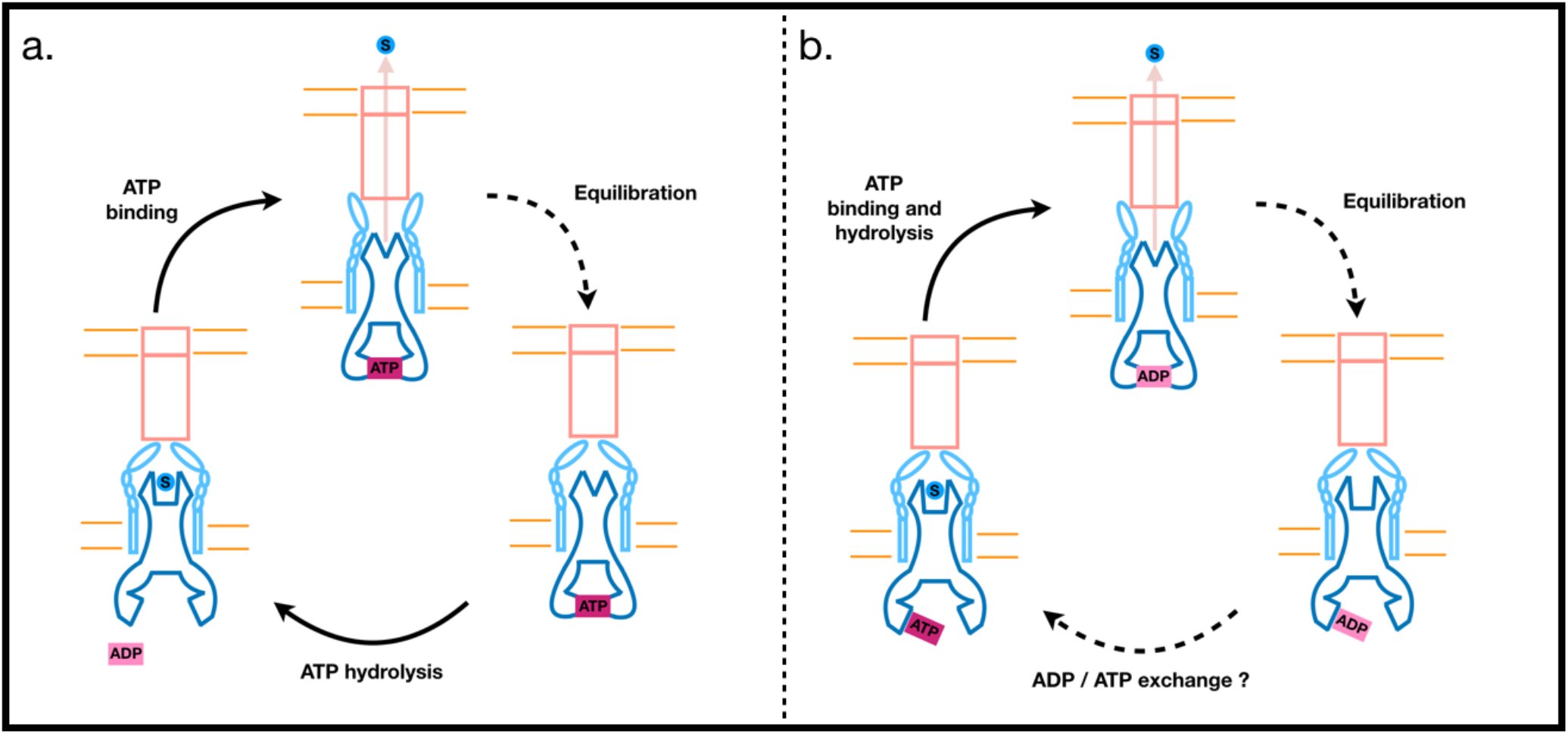
Two propositions of the molecular bellows mechanism for substrate secretion by the MacAB-TolC tripartite efflux pump. a) By analogy with a fireplace bellows, Crow et al. proposed a catalytic cycle^4^ where substrates are pumped out through TolC (orange) by mechanotransmission-induced changes in the periplasmic domains of MacB (dark blue) and prevented from flowing back into the periplasm by a valve in MacA (light blue). In this model, it is suggested that the powerstroke for transport occurs upon ATP binding and that ATP hydrolysis allows to reset the pump in the inward-open state. b) Based on our experimental data, we suggest a slightly modified description where the mechanotransmission is rather mediated by ATP binding and / or hydrolysis. The slow, rate-limited, resetting of the pump would be due to internal conformational changes, possibly associated with the release of ADP and re-binding of ATP.

However, our conclusions differ from that regarding the substeps described in the original fire bellow model^4^. Indeed, it was suggested therein that power stroke is due to mere ATP binding and that ATP hydrolysis is used to impose the directionality of transport. Our assay allows for the simultaneous monitoring of single-turnovers for both ATP hydrolysis and roxithromycin transport in real time and we show that both phenomena are biphasic and seemingly synchronous. By contrast to the model presented Figure 3a, we suggest that the first initial ATP burst is correlated to the actual transport event, triggered by the mechano-transmitted reduction of the periplasmic cavity volume that squeezes substrates upwards towards MacA and TolC. The slower phase is most probably due to further rate-limited events that could be the reopening of the periplasmic cavity upon relaxation of MacB stalk helices combined to conformational changes releasing the association between the NBDs. Altogether, this re-equilibration of the system allows for subsequent transport events to take place (Figure 3b). Lately, Robert Tampé and collaborators performed single-turnover assays for TmrAB in proteoliposome^28^ and concluded that ATP binding drives a stoichiometrically-coupled substrate translocation thanks to a conformational switch that reorients the protein from an inner facing to an outward facing conformation. We suggest that in the case of MacAB TolC, ATP hydrolysis is the power stroke for transport and that ATP binding would be the switch towards the re-priming of the pump.

In conclusion, our approach allows a quantitative, real-time analysis of a tripartite machinery. This methodology will probably be useful for the sensible interpretation of the increasing number of fascinating cryoEM and crystal structures of MacAB-like pumps published.

## Methods

### Materials and reagents

1-palmitoyl-2-oleoyl-*sn*-glycero-3-phosphocholine (POPC) was purchased from Avanti Polar Lipids (USA), C12E8 (Octaethylene glycol monododecyl ether, ≥98% purity, ref. P8925), Triton X-100 (BioXtra, ref. 9284), ATP (Adenosine 5’-triphosphate disodium salt hydrate, ref. A2383), ADP (Adenosine 5’-diphosphate ≥95%, ref. 01905), L-Lactic Dehydrogenase from rabbit muscle (ref. L1254), NADH (β-Nicotinamide adenine dinucleotide, reduced disodium salt, ref. N0786), Pyruvate Kinase from rabbit muscle (ref. P7768), PEP (Phospho(enol)pyruvic acid tri(cyclohexylammonium) salt, ref. P7252), roxithromycin (ref. R4394) and quantum dots (CdTe core-type, ref. 777935) were purchased from Sigma, SM2 Bio-beads were obtained from Bio-Rad. Ni Sepharose High Performance (His Trap HP) and Superose 6 10/300 column were purchased from GE Healthcare.

### Protein preparation

MacA, MacB and TolC were expressed and purified as previously described in ref.^23^ with slight modifications. The genes were individually cloned into pET22a and expression was realized in C43 strains in 2YT media. Overexpressions were induced by adding 0.7 μM IPTG when cultures reached OD_600_ = 0.4 (overnight induction at 20°C). For solubilization, Triton X-100 was used for MacA and C12E8 for MacB and TolC. Membranes were diluted in 20 mM Tris pH 8, 300 mM NaCl, 1 mM PMSF at 2 mg/ml (as determined by BCA) and detergent was added at 4°C under gentle stirring at a 1:2 w/w ratio for 1h for MacB and TolC or at a 1:10 w/w ratio for 2-3h for MacA.

The suspension was cleared by ultracentrifugation (1h at 100,000 g) and the solubilisate, to which 10 mM imidazole was added, was loaded at a flowrate of 0.5 mL/min onto a 5 mL Ni-NTA superflow resin column equilibrated with 20mM Tris pH 8, 250mM NaCl, 10% glycerol, 0.2% C12E8 or 0.2% Tx-100. During the wash step, detergent was progressively adjusted/exchanged to 0.03% C12E8 for all proteins. Elution was performed using increasing Imidazole steps and proteins were eventually exchanged against Tris pH 8 20mM, NaCl 150mM, C12E8 0.03% on desalting columns (PD-10).

### Reconstitution of MacAB in nanodiscs

POPC lipids were dissolved in methanol/chloroform (v/v), dried onto a glass tube under steady flow of nitrogen and followed by exposure to vacuum for 3 hours. The lipid film was suspended at 10mM in the reconstitution buffer (Tris pH 8 20mM, NaCl 150mM, C12E8 0.03% or Tx 0.2% and Glycerol 20% (w/v) and subjected to 5 rounds of sonication for 30 seconds each.

Proteins in 20 mM Tris pH 8, 250mM NaCl, 10% glycerol, 0.03% C12E8 were inserted into nanodiscs according to a previously established protocol^23^ with the following modifications. MacB in C12E8 was mixed with POPC and MSP (MSPD1) at a final 2.5:90:1 MSP:lipid:MacB molar ratio. MacB and MacA were preincubated for 1h at a 1:6 molar ratio and then mixed with POPC and MSP (MSPD1E3) at a final 2.5:90:1 MSP:lipid:MacA-MacB molar ratio. After 1h incubation, detergent was removed by the addition of SM2 Bio-beads (quantity equal to 30 x the mass of C12E8) into the mixture shaken for 1h at 4°C. Suspensions were then centrifuged 30 min at 50 000g and the supernant is used extemporaneously for immediate use.

### Reconstitution of TolC in proteoliposomes

The POPC film was suspended at 10mM in the same reconstitution buffer as above (Tris pH 8 20mM, NaCl 150mM, C12E8 0.03% and glycerol 20%) now supplemented with 20μM CdTe quantum dots. The suspension was heated for 10 min at 37 °C and was submitted to five freeze thaw cycles. The liposomes were extruded through 400-nm membranes and through 200-nm membranes. C12E8 was then added to reach a 1:1 detergent/lipid ratio (w/w). Proteins were added to the solubilized liposome suspension at a lipids/OprM 1:20 ratio (w/w). Detergent removal was achieved using Bio-beads at a Bio-bead/detergent ratio of 30 (w/w) at 40°C overnight. Untrapped quantum dots were removed on PD-10 columns.

### Protein quantification

After purification and reconstitution into nanodiscs, samples were loaded on 12% acrylamide SDS-PAGE gels without boiling. After electrophoresis, the gels were stained with Coomassie Blue and digitally scanned. Densitometry was performed using ImageJ software. The linear regions in the densitometry profile were determined by measuring the density of standards with known protein amounts as shown in Supplementary Figures S1 and S2.

### Measurement of the ATPase activity

ATPase activity was determined at 20 °C with a coupled enzyme assay in a PTI International type C60/C-60 SE fluorometer, with the excitation and emission slit widths set at 5 nm. The excitation wavelength was set at 350 nm and emission at 460 nm. Measurements were performed in a medium comprised of 20 mM Tris pH 8, 50 mM NaCl, 2mM MgCl2, supplemented with 5 mM MgATP, 0.1 mg/mL pyruvate kinase (75U/ml), 1 mM phosphoenolpyruvate, 0.1 mg/mL lactate dehydrogenase (150U/ml), and an initial concentration of about 10 μM NADH.

### Measurement of the Quantum dot fluorescence quenching

Equilibrium fluorescence experiments were performed with a PTI International type C60/C-60 SE fluorometer with constant stirring of the temperature-controlled cell (20°C) and with the excitation and emission slit widths set at 5 nm. The excitation wavelength was set at 350 nm and emission at 525 nm. Measurements were performed in 20 mM Tris pH 8, 50 mM NaCl, 2mM MgCl_2_.

## Acknowledgements

Laurent Catoire for help in the nanodisc reconstitution. Dmitry Shvarev, Cédric Orelle and Gregory Boël for their careful reading and useful comments on the manuscript.

## Author contributions

H.S and MP designed the experiments. Q.C, H.S and D.P produced and purified the proteins. H.S and Q.C. produced proteins in nanodisc. H.S and W.B carried out fluorescence measurements. M.P. wrote the manuscript with inputs from all authors.

---

**Supplemental Material: Figure S1.**
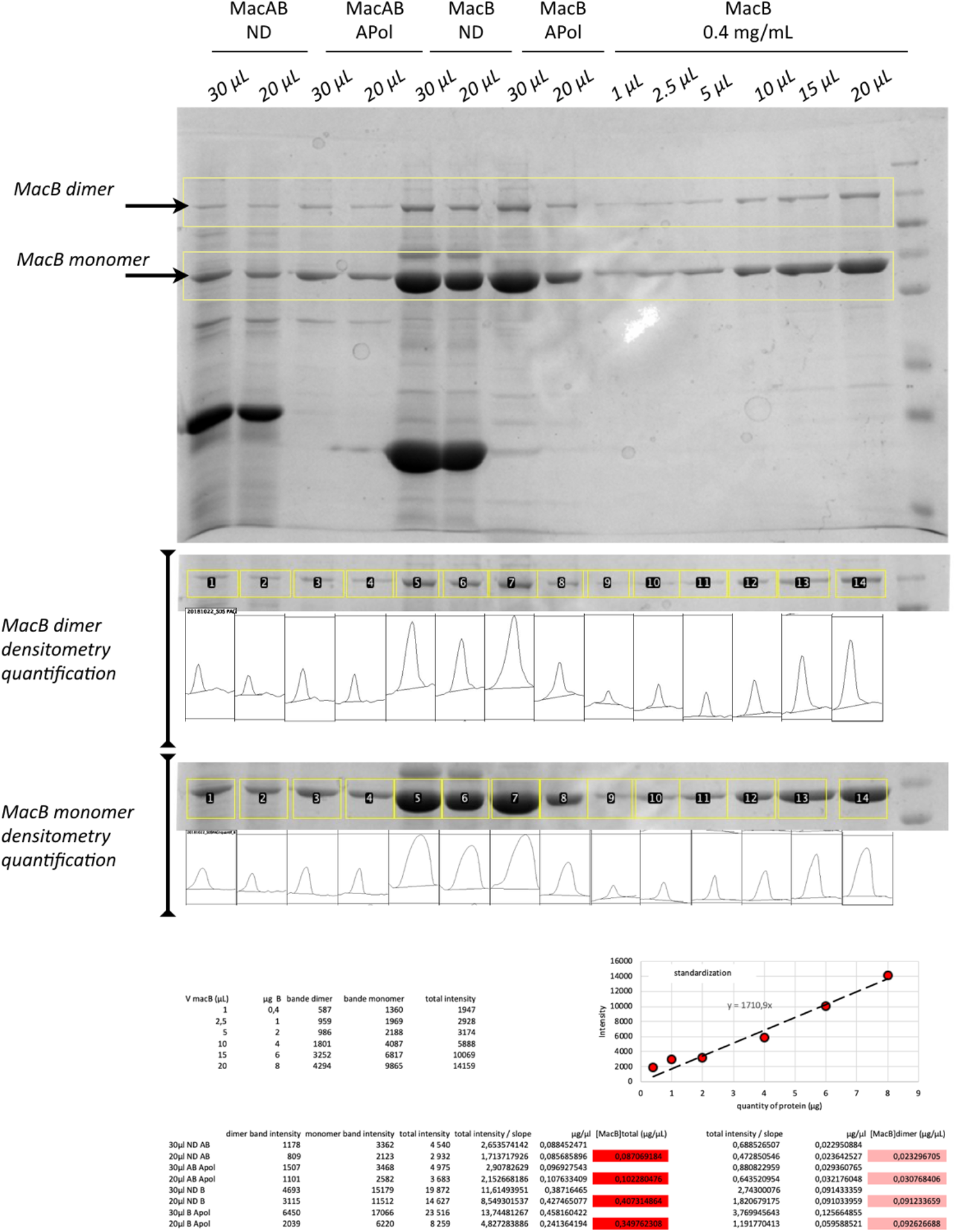
SDS-PAGE gel analysis and estimation of MacB quantity after purification and reconstitution into nanodiscs (ND) or amphipols (APol), prior to their use for the ATPase activity assays. Protein samples were diluted at a 1:1 ratio with Laemmli 2× buffer solution (Bio-rad) with 5% 2-mercaptoethanol (Sigma-Aldrich) as a denaturing agent and without heat denaturation. Protein samples (volumes indicated at the top of each gel) were loaded in the wells and electrophoresis was run at 200 V for 30 min in a Mini-Protean Tetra cell (Bio-rad) using TGX running buffer. Gels were washed with distilled water and stained with Coomassie Blue. and imaged in a Molecular Imager ChemiDoc XRS System (170-8070, Bio-rad) under white light epi-illumination. Images were saved as a TIFF file and analyzed using ImageJ (NIH). The Gel Analyzer tool of ImageJ was used to determine the profiles of each lane of the gel. The overall MacB protein load (monomer + dimer) or the proportion corresponding to MacB dimer only was correlated with the corresponding peak area in the densitometry profile (shown below each band selection, depicted by yellow boxes), shown at the bottom of the figure (standardization calculated in Excel spreadsheet).

**Supplemental Material: Figure S2.**
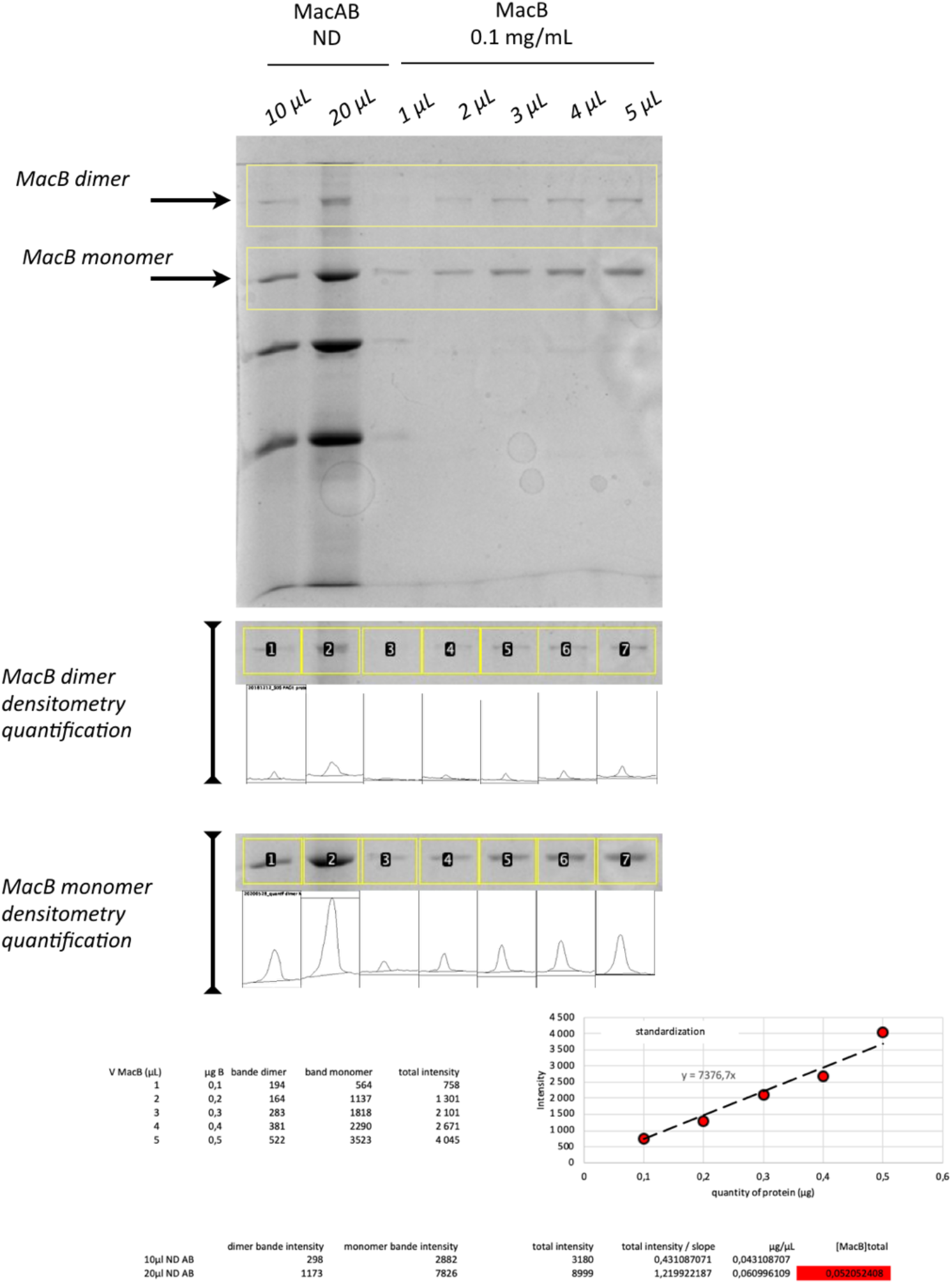
SDS-PAGE gel analysis and estimation of MacB quantity after purification and reconstitution into nanodiscs (ND), prior to their use for the roxithromycin transport assays. Protein samples were diluted at a 1:1 ratio with Laemmli 2× buffer solution (Bio-rad) with 5% 2-mercaptoethanol (Sigma-Aldrich) as a denaturing agent and without heat denaturation. Protein samples (volumes indicated at the top of each gel) were loaded in the wells and electrophoresis was run at 200 V for 30 min in a Mini-Protean Tetra cell (Bio-rad) using TGX running buffer. Gels were washed with distilled water and stained with Coomassie Blue. and imaged in a Molecular Imager ChemiDoc XRS System (170-8070, Bio-rad) under white light epi-illumination. Images were saved as a TIFF file and analyzed using ImageJ (NIH). The Gel Analyzer tool of ImageJ was used to determine the profiles of each lane of the gel. The overall MacB protein load (monomer + dimer) was correlated with the corresponding peak area in the densitometry profile (shown below each band selection, depicted by yellow boxes), shown at the bottom of the figure (standardization calculated in Excel spreadsheet).

**Supplemental Material: Figure S3.**
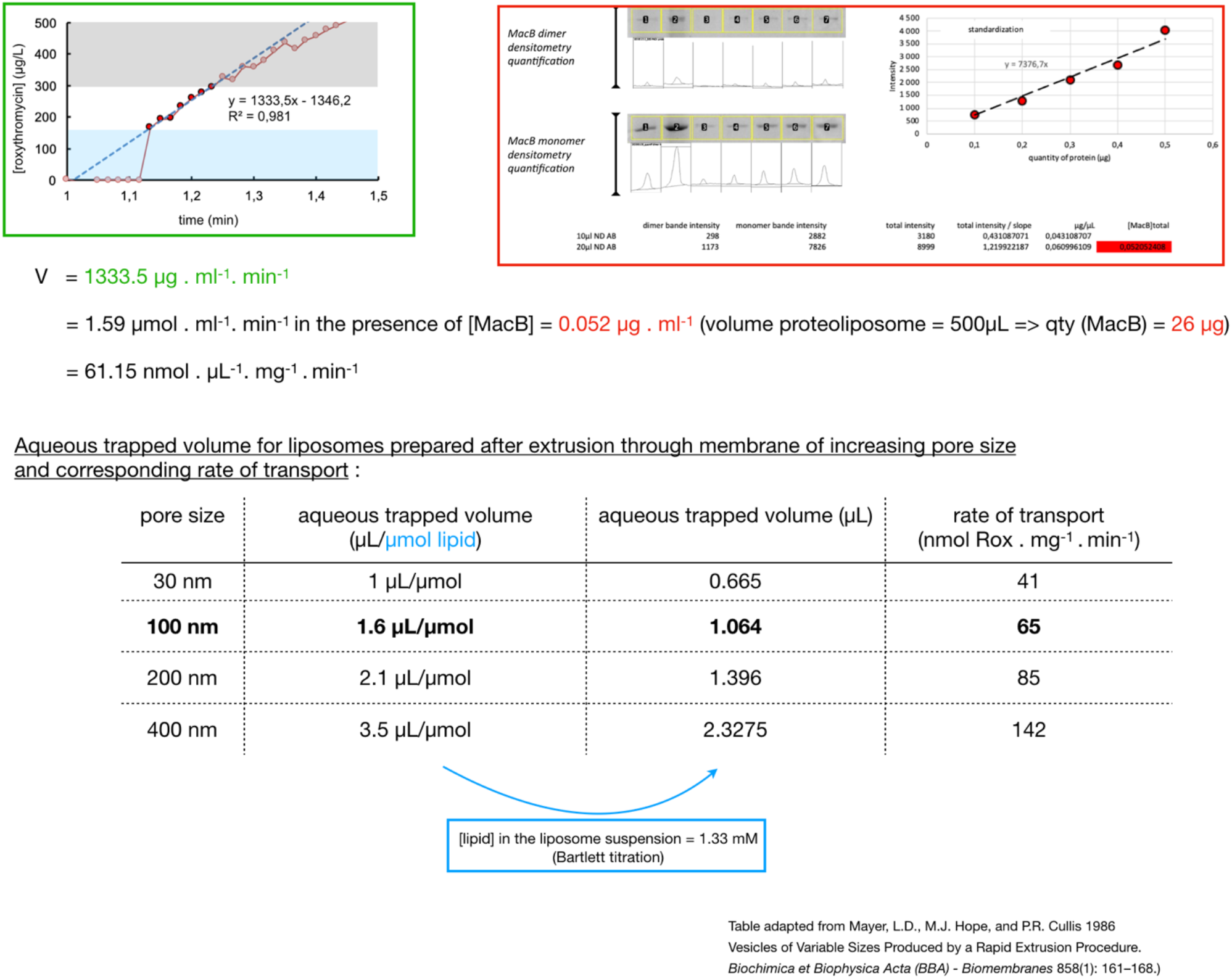
Details of the calculation. From the kinetics of roxithromycin transport (green panel reproduced from the experiment shown Figure 2b), we estimate a rate of steady-state roxithromycin transport of 1335.5 μg.ml^-1^ .min^-1^, corresponding to 1.59 μmol.ml^-1^ .min^-1^ (MW roxithromycin = 837 Da). From the densitometry measurements (red panel reproduced from the experiment shown Supplementary Figure S2), we estimate that this rate of transport was obtained from a quantity of 26 μg MacB in nanodiscs. Hence, we estimate a rate of 61.15 nmol of roxithromycin transported per mg of MacB, per minute in a volume of 1μL. [1] We evaluate the internal volume of the liposome by comparison with the work of Mayer and collaborators where they experimentally measured the aqueous trapped volume by titration of entrapped Na^22^ or [^14^C]inulin for liposomes prepared after extrusion through polycarbonate filters of increasing pore diameter. For liposomes prepared after extrusion through 100nm membranes, *i.e* in conditions similar to ours, they calculated a volume of 1.6μL per μmol lipid used for the liposome preparation. We calculated the lipid concentration by Bartlett titration and found a value of 1.33 mM. Hence we extrapolate that the aqueous trapped volume in our suspension is 1.064 μL. [2] From the above ([1] + [2]), we conclude that the rate of roxithromycin transport is 65 nmol per nmol MacB and per minute.

**Supplemental Material: Figure S4.**
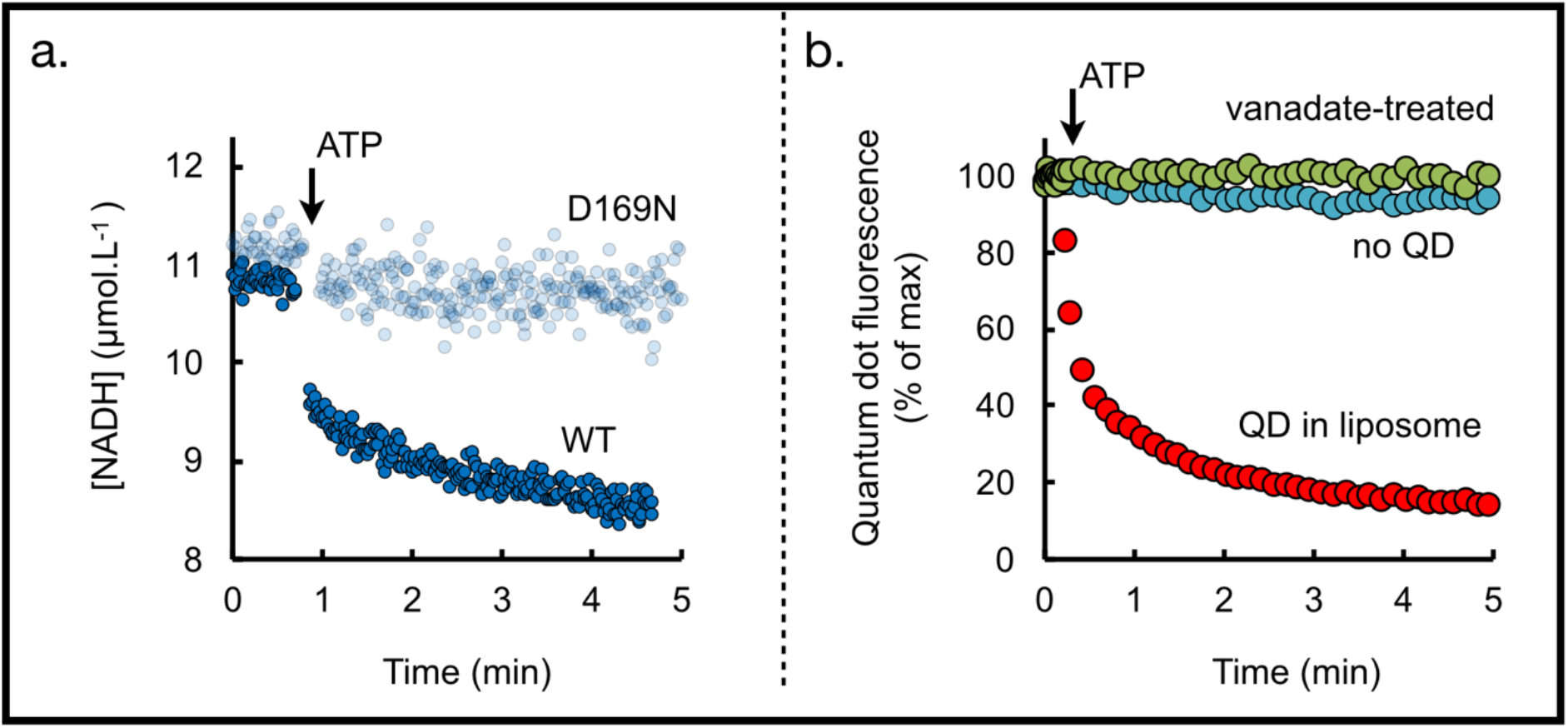
a) Variation of the NADH concentration as a function of time after 10-fold dilution of the MacAB nanodiscs (prepared from WT versus mutant MacB; *c.a* 10 μg MacB, as standardized by SDS PAGE densitometry, see supplemental Figure S1) into 20 mM Tris pH 8, 50 mM NaCl, 2mM MgCl_2_ containing the coupled-enzyme assay (see Methods for details) and addition of 1 mM ATP (represented by the arrow). The proportionality coefficient between NADH fluorescence and the NADH concentration allowed the conversion of the fluorescence changes (not shown here) into [NADH] variations. Changes due to dilution have been corrected for. NADH fluorescence was measured with an excitation wavelength set at 350 nm and emission at 460 nm. b) CdTe quantum dot fluorescence is measured as a function of time after 10-fold dilution of the MacAB / TolC complex into 20 mM Tris pH 8, 50 mM NaCl, 2mM MgCl_2_, addition of 7μM roxithromycin and 7μM vanadate (not shown here for the sake of clarity) and of 1 mM ATP (represented by the arrow). Measurements were performed in the presence of MacAB-ND and QD-loaded TolC proteoliposomes (red curve), in the presence of MacAB-ND and QD-free TolC proteoliposomes (blue curve) or in the presence MacAB-ND treated with vanadate and then mixed with QD-loaded TolC proteoliposomes (green curve). Vanadate acts as a potent inhibitor of many ATPases, because it can mimic the transition state for the γ-phosphate of ATP during hydrolysis and stabilize the transition state conformation. The MacAB nanodiscs (25μg MacB, as standardized by SDS PAGE densitometry, see supplemental Figure S2) and TolC proteoliposomes were preincubated for at least 1h at room temperature prior to their addition in the fluorescence cuvette, roxithromycin was added in the cuvette at least 10 minutes before addition of ATP, vanadate was added 1 minute after roxithromycin. Changes due to dilution have been corrected for. Traces were normalized as 100% right before addition of ATP.

